# Programmed ribosomal frameshifts, and how to find them

**DOI:** 10.1101/2023.04.10.536325

**Authors:** Katelyn McNair, Peter Salamon, Robert A. Edwards, Anca M. Segall

**Affiliations:** Computational Science Research Center, San Diego State University, San Diego, CA, USA; Department of Computational Science University of California Irvine, CA, USA; Department of Mathematics and Statistics, San Diego State University, San Diego, CA, USA; College of Science and Engineering, Flinders University, Bedford Park, Adelaide, SA, 5042, Australia; Department of Biology, San Diego State University, San Diego, CA, USA

## Abstract

One of the stranger phenomena that can occur during gene translation is where, as a ribosome reads along the mRNA, various cellular and molecular properties contribute to stalling the ribosome on a slippery sequence, shifting the ribosome into one of the other two alternate reading frames. The alternate frame has different codons, so different amino acids are added to the peptide chain, but more importantly, the original stop codon is no longer in-frame, so the ribosome can bypass the stop codon and continue to translate the codons past it. This produces a longer version of the protein, a fusion of the original *in-frame* amino acids, followed by all the alternate frame amino acids. There is currently no automated software to predict the occurrence of these programmed ribosomal frameshifts (PRF), and they are currently only identified by manual curation. Here we present the first machine-learning based method to detect and predict the presence of PRFs in all types of coding genes and taxa with an accuracy exceding 90%.

## Introduction

For the last half-century, the *Central Dogma* has shaped our understanding of molecular biology. In the idealized version of genetic information flow: DNA is transcribed into RNA, which is then translated into protein. During transcription, a polymerase reads along a strand of the DNA, one base at a time, catalyzing the production of a complementary strand of RNA. During translation, a ribosome reads along the RNA, one codon at a time, forming a polypeptide chain of amino acids until it reaches a *stop* codon. Since a codon is composed of a nucleotide triplet, the ribosome shifts three nucleotides at a time in order to maintain the fidelity of the amino-acid encodings. The first exception to this +3 indexing was hypothesized in 1975 to explain the occurrence of peptides slightly larger than expected during *in vitro* translation of the four known genes in the phage *MS2* genome. A small fraction (5%) of the synthetase gene, whose product is 62kDa, migrated during electrophoresis as if it were 66 kilodaltons, approximately 40 amino acids longer than expected (1). Stop codon readthrough was ruled out, and several possibilities were hypothesized: protein splicing, base insertion or deletions during transcription, post-translational modifications, and even that the ribosome might “retranslate” portions of the gene. However, it was the very last possibility surmised *“that 7 % of the ribosomes translating the synthetase shift out of phase and bypass the normal termination signals”* proved to be correct. In subsequent work, it was shown that the rate at which the larger protein was translated could be affected by adjusting the concentrations of the different tRNAs, and also revealed that one of the other four genes, a coat protein, also had variable protein sizes (2). Although the significance of the frameshifted synthetase product for phage replication has yet to be determined, the function of the frameshift in the coat gene was to terminate coat protein synthesis early and present the frameshifted ribosome at the start of the overlapping lysis gene (3). Presumably, once enough copies of the coat protein were translated, sufficient complete virions have been assembled, and sufficient levels of lysis protein have increased, it is then time to rupture and escape from the doomed host cell.

It was not until genetic sequencing became more widely available and the region around the known frameshift in the *gag*/*pol* gene of the mouse mammary tumor retrovirus *Rous sarcoma* was sequenced that a more detailed model of the necessary “signals” involved in backward ribosomal frameshifting was proposed. In this model, there is a slippery sequence upon which the ribosome shifts several bases, paired with downstream stem-loop or pseudoknot secondary structures, “*that may act by stalling translating ribosomes, thereby promoting the tRNA slippage*” of the bound codon:anticodon pairs in the E and P sites of the ribosome (4).

Over the years, other signals contributing to the backward model were identified, most notably ribosomal binding site (RBS) interference. Although first shown to play a role in forward frameshifts (5), it was later found that RBS interference also plays a role in backward frameshifting (6). Typically, bacterial translation initiation requires a Shine-Dalgarno (SD)-like sequence a few bases upstream of the start codon. This sequence promotes recruitment of the ribosome onto the mRNA because the small subunit of the 16S rRNA molecule has a complementary *anti* Shine-Dalgarno sequence found near the entrance channel of the ribosome that recognizes and binds to the RBS. Therefore if during translation, the ribosome encounters a slippery sequence that has an RBS-like motif properly positioned on the mRNA, it interacts with the *anti* Shine-Dalgarno sequence on the ribosome and facilitates frameshifting.

Surprisingly, the only slippery sequence motif identified to date is the original heptamer motif XXXYYYZ, first posited for the *gag*/*pol* gene backward frameshift (7). This mechanism involves the simultaneous (backward) slippage of two tRNAs along the mRNA within the decoding center of the ribosome (Figure 1). During translation, when the ribosome reaches the slippery site and the two tRNAs XXY and YYZ are bound to their respective codons on the mRNA, cellular signals combine to shift the two tRNAs backwards one base, so they are paired with XXX and YYY codons (Figure 1). Even though the new pairing is not exact, since each codon/anti-codon pairing has one mismatched nucleotide, the wobble nature of the third position allows for transitory stability until the ribosome recruits a tRNA matching the new -1 A-site codon, and translation continues in the -1 frame. For forward frameshifts, no such general motif has been identified yet. Instead, a ribosome encountering an unfulfilled rare codon during a gene translation can pause long enough to cause a forward shift into the +1 frame. It is generally thought that 3’ secondary structures do not play a role in +1 forward frameshifts; however, a 3’ pseudoknot plays a role in the +1 programmed ribosomal frameshift of mammalian ornithine decarboxylase antizyme (8). The presence of such knots downstream of +1 forward frameshift sites might instead be the result of bidirectional ribosomal frameshifting, which has been shown to occur in both the human *prfB* gene (9) and the ORF1a polyprotein of the SARS-Cov-2 coronavirus (10). In the bidirectional model, secondary structures on both sides of a slippery site help to regulate ribosomal frameshifting; secondary structure on one side acting to nudge the ribosome backwards or forwards, while an attenuator structure on the other side works to block the shift. The structure downstream of the slippery sequence can be either simple hairpins or more complex pseudoknots, while upstream structures are limited to hairpins since they are quicker to form once the ribosome has passed. The full extent of bidirectional control at slippery sites has yet to be determined; perhaps all frameshifts have some limited bidirectional control, and we have only identified the more obvious antipodal pseudoknots, while ignoring the lesser hairpins on the attenuation (upstream) side.

**Figure 1).**
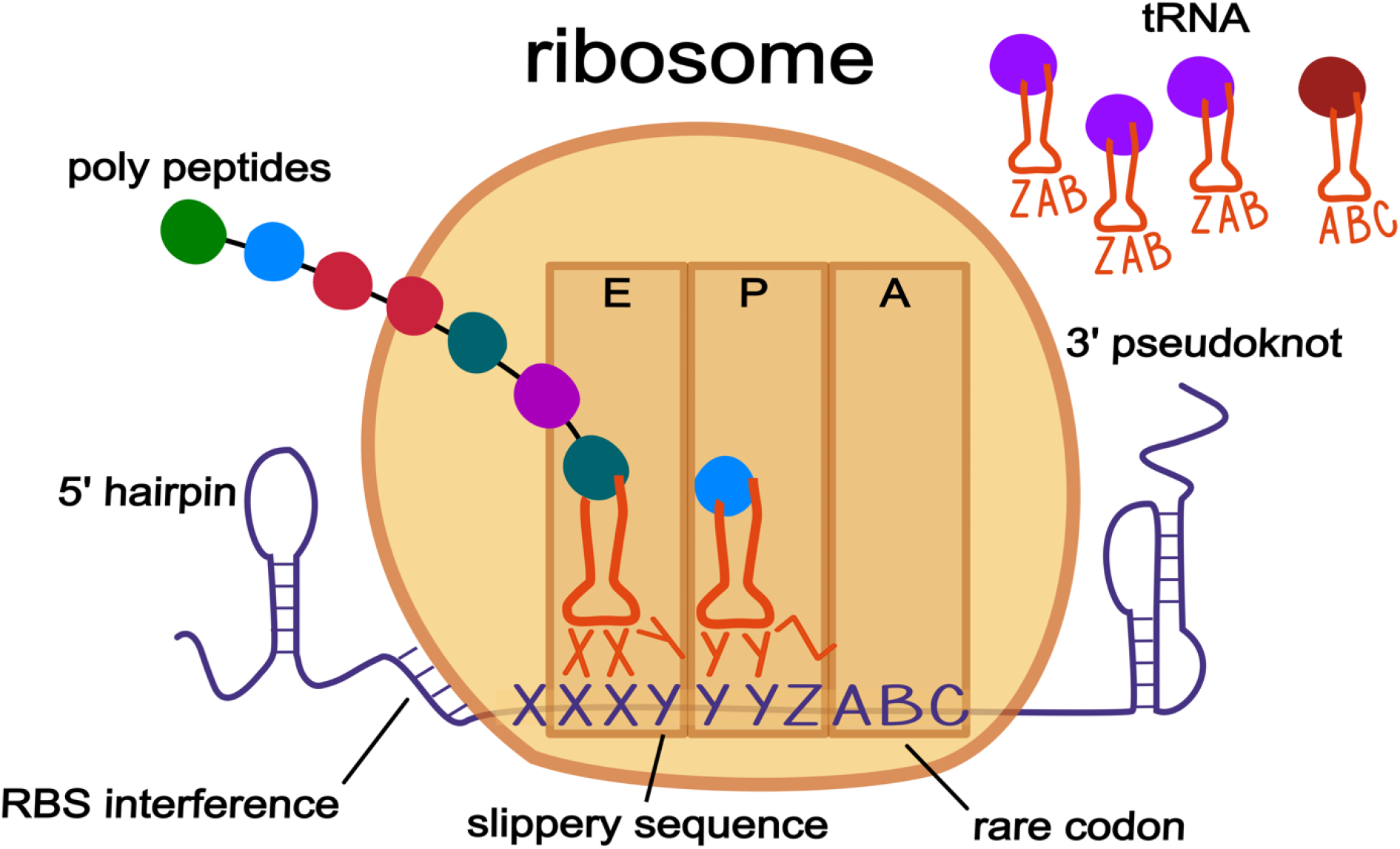
Some known cellular properties that are thought to contribute to programmed ribosomal frameshifts. The bi-directional model is shown with backwards -1 frameshifts, although the properties may also be associated with forward +1 frameshifts. The general model is based around a slippery sequence, in this case the canonical XXXYYYZ motif that was first identified in the gag/pol gene of a murine retrovirus. During translation as the ribosome is reading along in the 5’ to 3’ direction, it encounters the slippery site in its E and P site, which are both filled with their matching cognate tRNA. The presence of 3’ secondary structure along with a 5’ ribosomal binding like sequence work to pause the ribosome. The presence of a rare codon (ABC) is generally thought to induce forward frameshifts, however the ratio of the waiting A-site codon (A0) versus the ratio of the codon shifted into (A1) may also be involved in backwards -1 frameshifts since moving the A-site into frame with a rare codon would be unfavorable. While paused at the slippery sequence, the absence of 5’ secondary structure allows the ribosome to slip backwards 1nt putting it into the -1 frame. The motif of the slippery site allows the anti-codons of the tRNAs (XXY and YYZ) to satisfactorily re-bind with the -1 codons (XXX and YYY) due to the permissible nature of the third—wobble—position. The new A-site codon (ZAB) is then filled with its cognate tRNA, the bases pairing of the 3’ secondary structure momentarily disassociates, allowing the ribosome to proceed with translation in the -1 frame.

The lack of experimental evidence for slippery site motifs other than the original XXXYYY sequence as well as limited knowledge of the full extent of cellular properties that contribute to inducing ribosomal frameshifts, makes creating software to predict their occurrence in genomic data rather difficult. This might account for why there is not a single universal tool available for predicting programmed ribosomal frameshifts. The few available tools that investigate programmed ribosomal frameshifts do not leverage machine learning methods since they only look for a single specific slippery sequence motif and show all possible locations of that motif (11,12), or were designed to only predict frameshifts for a specific gene of a specific taxon (13,14). Here we present PRFect, a new computational tool that is intended to predict ribosomal frameshifting of all types of coding genes in complete genomes from all domains of life, that is both accurate and also very easy to use.

## Methods

### github.com/deprekate/prfect

All the code and data presented here are available in the GitHub repository. All of the code exclusive to the PRFect package was written in Python3 in order to be user-friendly and easily updateable for future improvements. PRFect is also available on the Python Package Index PyPI (pypi.org) as an easily installable command-line program that downloads and installs with a single command: *pip install prfect*. The PRFect package does require the third-party dependencies: *scikit-learn* and *numpy* (*15,16*), as well as the additional packages, *genbank*, *score_rbs*, *linearfold*, and *hotknots.* The last two were adapted from their original C code libraries (17,18) into Python packages that auto-install along with the other packages when the previous command is used to install PRFect.

### Obtaining Data

To obtain ribosomal frameshift data, we downloaded 3,679 phage genomes in GenBank format from the Actinobacteriophage Database phagesdb.org (19). Genes exist as *CDS* features within the GenBank format (20) and no explicit designation indicates if or where ribosomal frameshifting occurs within a gene. However, when a *CDS* feature has discontinuous locations in the GenBank file, they are denoted by using the *join* keyword in the coordinates. Figure 2 shows a small example GenBank file with two genes. The first example gene occurs from nucleotide 1 to nucleotide 100, while the second example gene exists in two locations, which are also two different frames, through the use of the *join* keyword.

**Figure 2).**
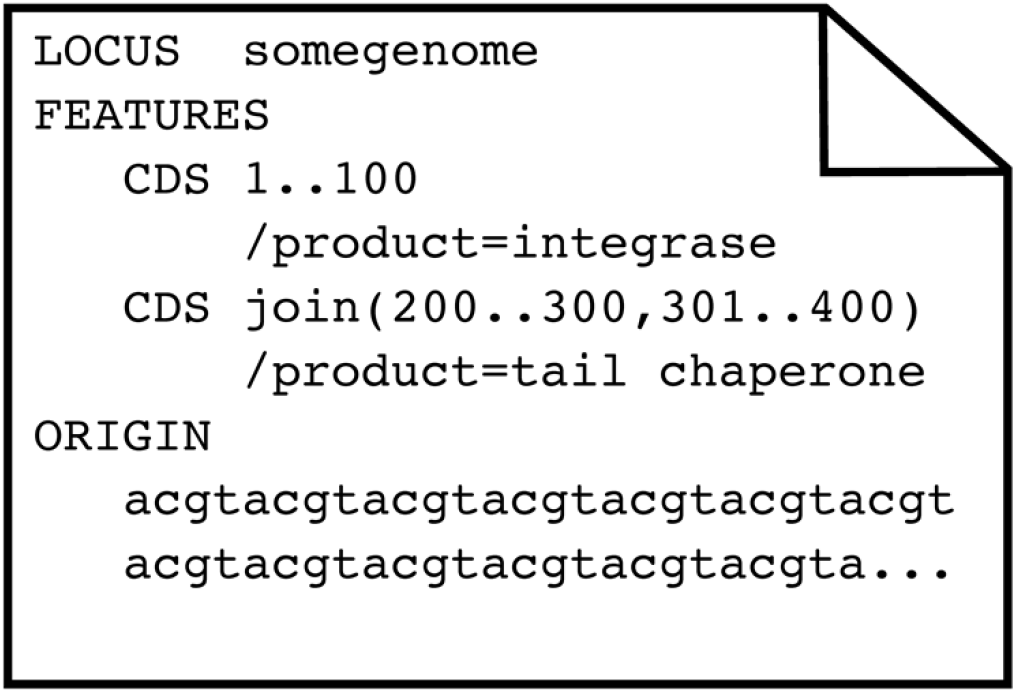
A graphical representation of an exemplar GenBank file. A GenBank file has various keywords in a specific order. Only the three most pertinent sections are shown: LOCUS, FEATURES, and ORIGIN. The first line of the file always contains the keyword LOCUS followed by the name of the genome. Then comes the keyword FEATURES followed by lines that contain the type of feature, in this case, coding sequences (CDS) and the location within the genome of that feature. The features can have descriptor tags that describe the various properties of that feature. Each feature tag begins with a forward slash, and in this case the two features are tagged with functional annotations through the use of the /product keyword. The last important section is the ORIGIN, which contains the DNA or RNA backbone of the respective genome. More details about the GenBank flat file format can are on the NCBI website (https://www.ncbi.nlm.nih.gov/Sitemap/samplerecord.html).

Since frameshifted genes have multiple locations, they can be found by looking for CDS *Features* that use the *join* keyword. However, in addition to ribosomal frameshifting, other reasons can cause a gene to exist in multiple locations, such as splicing, mobile elements that insert into genes, genes spanning break points in circular genomes, and even sequencing errors. To distinguish ribosomal frameshifted genes from other causes, we required only two sets of coordinates within 10 bp of each other, which was chosen through manual inspection of the genomes. Ideally, the two pairs of ribosomal frameshifted genes would be separated by either 0 (backward) or 1 (forward) nucleotides. However, the *joined* ribosomally frameshifted genes of the SEA-PHAGES data varied from 0 to 7 nucleotides apart, while the genes that were *joined* due to other causes (discussed below) varied from 50 to 818 nucleotides apart. Of the 3,679 genomes, 2,489 phages had one or more *CDS* features with the *join* keyword, giving a total of 2,557 *joined* CDS *Features*. Of these, 61 were at opposite ends of the genome, indicating genes split due to the circularization of the genome. There were 20 *joined* features with coordinates separated by more than 10 bp that were excluded from the training data: six in genes coding for major capsids, lysins, and DNA methylases (caused by inteins from homing endonucleases); twelve in genes coding for minor tail proteins (due to group 1 introns); one encoding a “structural” protein; and one encoding a tail assembly chaperone. The two pieces of the tail assembly chaperone were separated by 72 bp, which we predict was an annotation error, along with all the other joined genes with coordinates further than 10 bp apart. There were 2,476 frameshifted genes (one per genome) and 360,977 genes without frameshifts. The frameshifted genes were split into their two constituent fragments, giving sets of two consecutive “pseudo” gene pairs for positive cases. For example, the frameshifted chaperone gene in Figure 2 was split into two CDS *features* with locations (200..300) and (301..400), and the *join* keyword was removed. Then all of the genes were evaluated pairwise to determine if an overlap could occur between consecutive genes, allowing for the possibility that the two genes are a single frameshifted gene. Figure 3 shows an example of such an overlap in a pair of consecutive genes (denoted in green).

**Figure 3).**
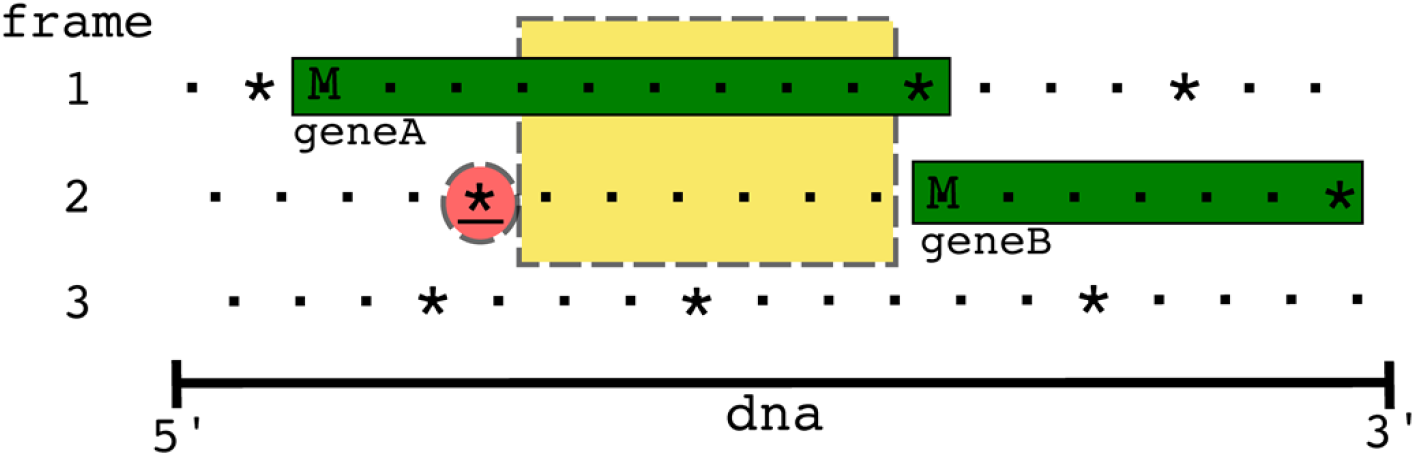
An example genome with two overlapping genes. Dots represent codons, asterisks represent stop codons, and M represents start codons. Only the forward three frames are shown. There is a gene in the first frame whose stop codon overlaps with the start codon of the gene in the second frame, however since the closest (red underlined) stop codon in the 5’ left direction from the start codon of *geneB* is six codons away, there is the possibility of a frameshift occurring in this overlap (yellow dashed box) region. If there were a frameshift that occurred, *geneB* would not be a complete gene but rather part of an alternate frameshifted translation of the fusion *geneAB*. The ribosomes that are frameshifted during translation end up producing this *geneAB* within the cell while the ribosomes that are not frameshifted only produce *geneA*.

In the example, *geneA* (frame 1) is 10 codons long and overlaps with 1 codon of *geneB*; since the first stop codon upstream (in the 5’ direction) from *geneB* (frame 2) is six codons to the left, there is the possibility that a frameshift could occur within this overlap region (yellow in Figure 3). If there were a frameshift that occurred, *geneB* would not be a complete gene but rather part of an alternate frameshifted translation of the fusion *geneAB.* During translation the ribosomes that are not frameshifted end up producing *geneA* within the cell while the ribosomes that are frameshifted end up producing *geneAB.* Most consecutive genes were either on opposite strands (thus, ribosomal frameshifting is untenable) or did not have the possibility of overlap, leaving only 67,664 gene pairs to search for negative case slippery sites.

### Slippery Site Motifs

The only slippery sequence to appear in the literature is the previously mentioned XXXYYY motif (7). However this “*threethree* motif” (the number denotes the same nucleotide repeated three times and then another nucleotide repeated three times) is not present in most frameshift overlaps, so we explored recurring patterns that could similarly serve as motifs. Originally, we tried using various automated tools to detect motifs, such as the MEME Suite (21), but this proved troublesome due to the highly repetitive nature of our training data, so we were forced to manually inspect the overlap regions of the annotated frameshifts. Focusing only on those annotated frameshifts that lacked the *threethree* motif, we looked for novel nucleotide patterns that could function similarly to the *threethree* wobble base pairing dynamic. For backwards frameshifts, we found the eight different slippery sequence motifs: *six*, *threethree*, *fivetwo*, *twofive*, *twofour*, *threetwotwo*, *five*, and *twoonefour*. A description of these motifs is in Supp Fig 1. For forward frameshifts we only looked for the two motifs *four* and *three;* however, we also required that the codon of the +1 frame A-site relative abundance (A1) is greater than the codon of the +0 frame (A0) A-site. Requiring the A1 codon relative abundance to be more favorable limits the number of candidate forward slippery sites found and speeds up runtimes because three (and four) bases in a row occur very often in a genome. Since some motifs are subsets of other motifs (i.e. *twofive* is also *twofour*, *six* is also *threethree,* and *four* is also *three*), the motif with a lower probability of occurring randomly takes precedence. Once all possible motifs were identified, we were left with a set of 106,692 different sites as potential slippery sequences, of which 3,711 were from (*true*) frameshifted genes, and 102,981 were from not thought to be frameshifted gene pairs (*false*). Since the frameshifts are not experimentally tested, we do not know the actual location of the slippery site, only the approximate location that the researcher guessed it to be. Therefore we cannot ascertain exactly which of the motifs in a *true* frameshift overlap region is the actual slippery sequence. To mitigate error induced by including incorrect, randomly occurring slippery sites into the dataset as *true* cases, any such motifs that occurred further than 10bp away from where the shift was annotated to occur were denoted as *false* cases. This left 2,718 *positive* cases (2,368 backward and 350 forward) and 103,989 *negative* cases.

### Properties contributing to translation efficiency (and potential pausing)

For every slippery sequence motif, cellular properties relevant to the translation process were aggregated based on the motif occupying the E-site and P-site of the ribosome, with the A-site being empty and waiting for tRNA recruitment to occur. The first property was the direction of the frameshift, forward or backward. The next two properties were the relative frequency of the waiting +0 A-site codon, and the relative frequency of the -1 or +1 frameshifted A-site codon. The frequencies are found during the genome file reading step by iterating through all the annotated coding genes and counting the relative occurrence of the 64 different codons. This accounts for the idea that if the codon waiting in the +0 A-site has few matching cognate tRNAs available in the cytoplasm while the tRNA for the ±1 A-site codon is quite abundant, the A1 codon is slightly more favorable than the A0 codon, and so the occurrence of a frameshift is more favorable. The next two properties added were different methods for scoring the presence of a ribosomal binding site (RBS) upstream of the P-site. The first method is a reimplementation of the 28 variable bins utilized by Prodigal for gene calling, where each bin is an integer from 0 to 27 and corresponds to a given RBS motif and nucleotide spacer sequence (22). The other is a reimplementation of the method employed by the RAST website, which uses the observed frequencies of 191 different RBS motifs with 10 different nucleotide spacer sequence sizes (23). To estimate the 3’ secondary structure, the minimum free energy (MFE) of the 50 bp and 100 bp windows were added using two different secondary structure prediction tools: LinearFold, which predicts simple hairpins, and HotKnots, which can predict pseudoknots. LinearFold produces MFE prediction scores identical to the widely used (but not available as an easily installable Python package) RNAFold program from the Vienna software package (24). For more complex secondary structure predictions that include pseudoknots, the HotKnots tool was run with the most recent DP09 parameter set. A parameter sweep of the data using pairs of window sizes from 30 bp to 120 bp, in 10 bp increments, and with offsets of 0 to 6, in 3 bp increments was attempted. The results were ambiguous, and no apparent accuracy peaks were observed, so we used the conventional 50 bp and 100 bp windows taken just after the three A-site bases of the slippery sequence (an offset of 3). The MFE scores were scaled by dividing by the window length, and since the MFE score is biased by the GC content of the window in question (25), we further normalized the MFE/bp by also dividing by the GC content. A visual representation of this transformation is shown in Supp Figure 2.

The last property added to the model was the number of bases between the slippery sequence and the +0 *in-frame* stop codon. This property helps distinguish between more probable motifs near the *in-frame* stop codon from those that occur randomly much further upstream of the *in-frame* stop codon. For instance, in the example in Figure 3, slippery sites that occur towards the right side of the yellow box would be slightly more probable, or at least differentiable, than those occurring towards the left side of the box.

**Table 1.** The various cellular properties used to classify slippery sequences

| property | description | range |
| --- | --- | --- |
| **DIR** | The direction of the frameshift | -1, +1 |
| **RBS1** | The Prodigal ribosomal binding site score | 0 - 27 |
| **RBS2** | The RAST ribosomal binding site score | 0.0 - 6.3 |
| **MOTIF** | The slippery sequence motif | 0 - 9 |
| **A0** | The frequency of the A-site codon usage in all genes | 0.0 - 1.0 |
| **A1** | The frequency of the +1 A-site codon usage in all genes | 0.0 - 1.0 |
| **LF50** | The normalized LinearFold minimum free energy calculation of the 50bp window | 0.0 - 1.0 |
| **LF100** | The normalized LinearFold minimum free energy calculation of the 100bp window | 0.0 - 1.0 |
| **HK50** | The normalized HotKnots minimum free energy calculation of the 50bp window | 0.0 - 1.0 |
| **HK100** | The normalized HotKnots minimum free energy calculation of the 100bp window | 0.0 - 1.0 |
| **N** | The distance of the slippery sequence from the *in-frame* (+0) stop codon | 0 - ∞ |

These eleven (DIR, RBS1, RBS2, MOTIF, A0, A1, LF50, LF100, HK50, HK100, N) properties were then used to train a histogram-based gradient boosting classification tree to predict the direction of the frameshift: 0 for no frameshift, -1 for backwards, and +1 for forward. Since only -1 and +1 ribosomal frameshifts appear in the SEA-PHAGES data, and other frameshift size categories (such as -2 and +2) were omitted, as discussed further below. The HistGradientBoostingClassifier module from the Python Scikit-Learn package was used with default parameters except for the L2 regularization parameter, which was set to 1.0 to help prevent overtraining on the data. In addition, the *early_stopping* was turned off so that the training/validation/testing results would be deterministic rather than stochastic. Since multiple potential slippery sites can occur within the *overlap* region of a single pair of consecutive genes, only the highest-scoring slippery site (if any) is returned as the predicted PRF site by PRFect.

### Training and Validation Data

The SEA-PHAGES genomes are highly repetitive; many of the phages have near identical, closely related, taxa in the database, and some phages are exact duplications of other genomes. The presence of multiple copies of the same genome makes splitting the data into training and validation more complex; therefore, four different *leave-one-out* levels were used: CLUSTER, SUBCLUSTER, MASH95, and GENOME. CLUSTER and SUBCLUSTER are the two taxonomic levels that phages are assigned to during the SEA-PHAGES workflow (26). For example, in a CLUSTER split, all of the phage genomes of one CLUSTER are removed, a HistGradBoost model is trained on the remaining genomes, and then predictions are made for the genomes of the omitted CLUSTER. As there are 89 different CLUSTERS, 89 different validation models were built, and the resulting predictions were merged. Likewise, there are 102 SUBCLUSTERS, hence 102 models were built, and the predictions at that level were merged. Since there are so few CLUSTERS and SUBCLUSTERS represented in the data, and the lowest taxonomic level GENOME is dubious due to the training set contamination discussed above, an intermediary taxonomic level, MASH95, was calculated for all of the genomes. This taxonomic level was assigned by using the genome distance estimation of MASH (27) to cluster the genomes at 95% identity, which is analogous to a MinHash distance of 0.05. The parameters used were the same as recommended in the publication: a sketch *size* of 400 and *k* of 16.

### Testing Data

To assess the performance of *PRFect* on unrelated genomes and non-chaperone genes, a general PRF model was first trained on all of the SEA-PHAGES genomes that contained a *joined* tail assembly chaperone (TAC) gene, and then five different sources of frameshift data were tested. The first is the RECODE database (28); because it is currently unavailable, an archive.org snapshot of the 2010 downloadable files was used. The data includes many types of coding anomalies, spans all domains of life, and covers all gene functions. Each entry in the database corresponds to a single gene, for which the nucleotide sequence and site of the frameshift is given. From the SQL file, 725 entries labeled as “ribosomal_frameshift” were extracted and converted to GenBank files, each of which contained only a single gene. The second source of frameshift data was the set of 28 phage genomes listed with a potential frameshift site in the region analogous to the tail assembly chaperone gene of phage lambda (29). We downloaded the accessions that were listed in the supplementary documents from GenBank, and for those genomes that were missing the specified frameshift (as a joined CDS feature), we manually added the gene to the file at the specified slippery sequence location. The third source of frameshift data was the FSDB (Frameshift Database) which contains 253 frameshifts in a graphical website (30). The website was parsed to get the GenBank accession number of each frameshift, the slippery sequence, and the location of the slippery sequence, which was then used to retrieve the files from GenBank. In contrast to the other test datasets, due to the size and number of genomes in the FSDB an automated python script was used to add a *joined* feature to the GenBank file at the location specified in the frameshift data. Additionally, any pre-existing *joined* features were left in place. The fourth source of frameshift data was 106 virus genomes with known or predicted occurrences of ribosomal frameshifting in genes of different functions (31). The provided accession numbers were used to download the genome files from GenBank, and as before, frameshifted genes were added to those files lacking the specified feature. The fifth and last source of frameshift data was the quite topical single genome for the coronavirus Covid19. The GenBank file for sars-cov2 (accession NC_045512) contains 12 genes, one of which is frameshifted (32).

## Results

### Validation Sets

To estimate the accuracy of *PRFect*, a leave-one-out training validation approach was used at each of the four different taxonomic levels. At each level, the different groups of that level were iterated, leaving out all genomes of the iterated group, training a model on the remaining groups, predicting frameshifts on the left-out group, and then merging the predictions of all groups. Each potential slippery site either comes from a *joined* gene (i.e. a tail chaperone) or from two non-joined adjacent overlapping genes that happen to have a spuriously occurring slippery sequence motif. The PRFect algorithm was used to predict whether the potential slippery site promotes PRF or not (Supp Data). At the highest taxonomic validation split (leaving out all genomes of the same CLUSTER from training), out of 2,476 *joined* known PRF genes, 1,486 were correctly predicted as having PRF, which is a 61% recall (Figure 4, Seaphages Cluster).

**Figure 4).**
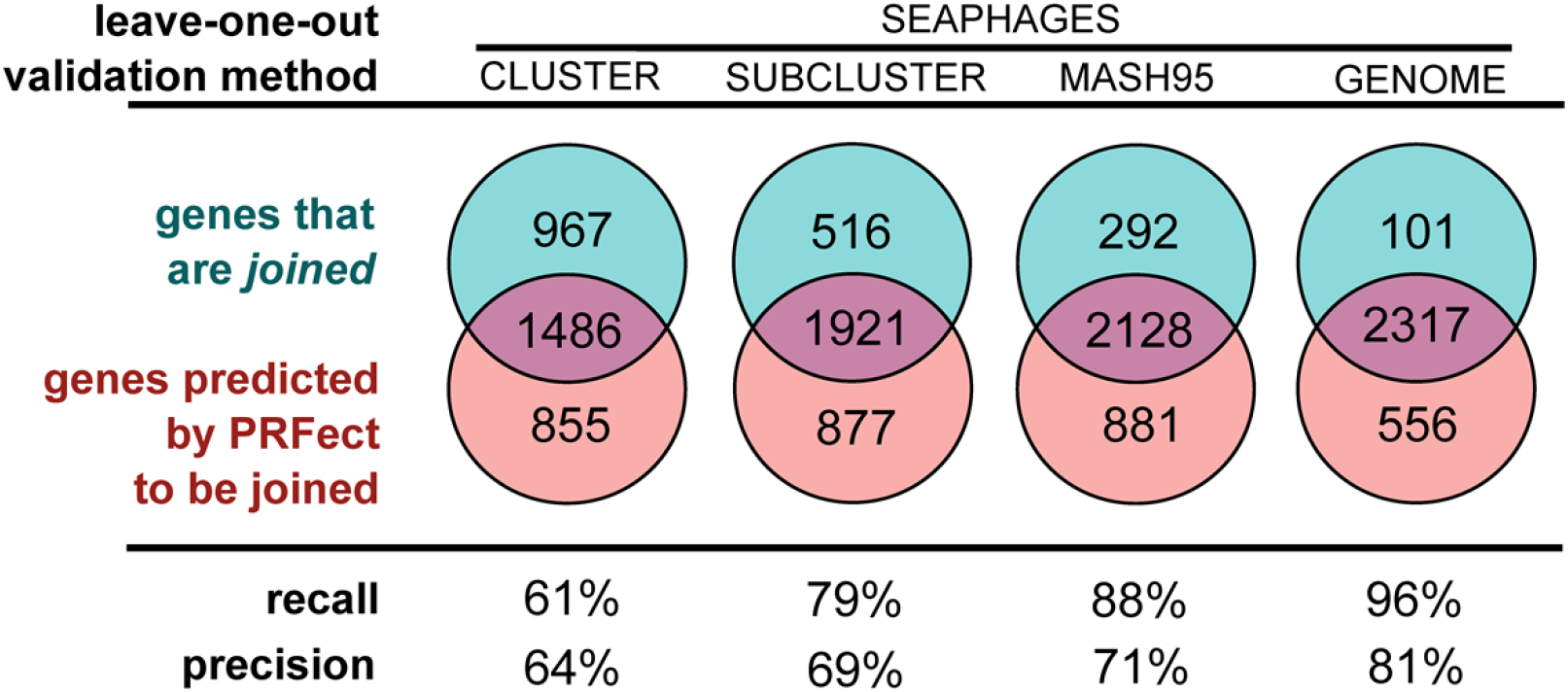
Training and leave-one-out validation on the SEAPHAGES data. While there were 3,679 genomes in the SEAPHAGES database, only 2,476 of them had a tail chaperone gene that was annotated as being frameshifted based on the gene having two locations in the genome linked by the use of the *join* keyword. A simple leave-one-out validation at the GENOME level could not reasonably be used alone to estimate the accuracy of PRFect, since the same genome—or a very close relative—might be present multiple times in the database. Likewise, the CLUSTER|SUBCLUSTER level validation was too broad, so an additional taxonomic level was created using the MINHASH algorithm, and genomes were clustered at 95% genome identity.

As expected, when the models are trained on more data, that includes similar genomes, the recall greatly increases, where at the lowest taxonomic validation split (leaving out only the single respective GENOME and then training on all other GENOMES) there were 2,317 out of the 2,476 *joined* known PRF genes predicted as frameshifting (True Positives), which is a 96% recall rate. As discussed in Methods (Training Data), since nearly the same genome may be found in the SEA-PHAGES data more than once, the GENOME level validation may be biased. Hence, the true real-world recall rate of PRFect falls somewhere between the CLUSTER level and GENOME level, depending on how similar a newly sequenced input genome is to previously known genomes used for training. Not shown in Figure 4 are all of the non-*joined* genes that were correctly predicted as not frameshifting (True Negatives), and only around 1,000 out of the 360,000 total non-*joined* genes in the SEAPHAGES genomes were incorrectly predicted as having a frameshift (Supp Figure 3), giving the models an *accuracy* of 99%.

### Testing Sets

Five alternate data sources were assembled to evaluate the performance of the pre-trained models on data not seen during training and different from the Actinobacteria phage genomes, which are known to have high %GC sequences. These five sources ranged from manually-curated online databases of frameshift data from all domains of life to lists of known frameshifts from publications to the single genome of the coronavirus SARS-CoV2 that causes the disease Covid19.

**Figure 5).**
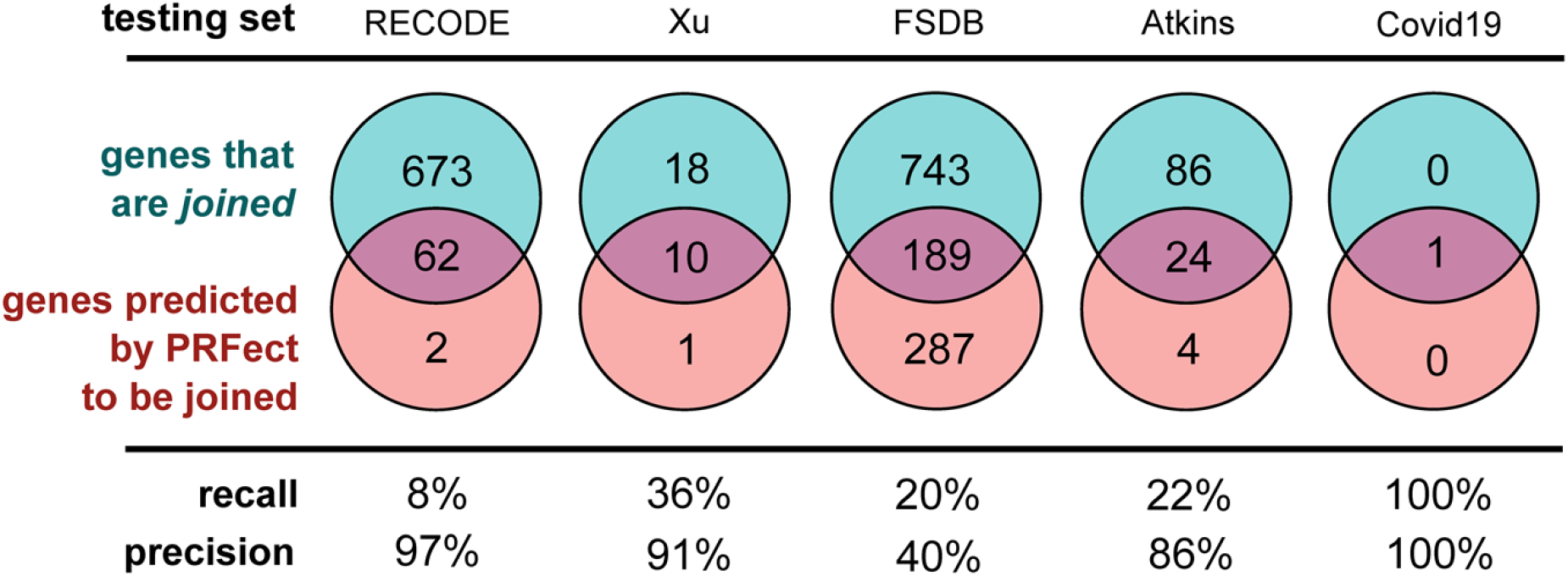
The various performance scores were calculated the same way as the previous validation accuracy, where the predicted slippery site had to belong to a *joined* gene and be within 10 bases of the frameshift annotation to be labeled a True Positive prediction. Predicted slippery sites further than 10 bp, or from genes that were not *joined*, were considered False Positives. True and False Negatives were labeled using a similar scheme.

### The RECODE dataset

The RECODE data contains 725 ribosomal frameshift sites, each of which is composed of only the single *joined* gene in question. The database contains 244 backward frameshifts and 481 forward frameshifts. As previously mentioned, the sequence files of the RECODE dataset are composed of only a single gene and not the entire genome. PRFect cannot adequately calculate codon usage to identify the rarity of each codon, which is why all of the 56 correct, true positive predictions for the RECODE data belong to backward frameshifts, while none of the 481 forward frameshifts was predicted correctly.

### The Xu phages

As mentioned, the RECODE data is not representative of real-world genomic data since it is only the single frameshifted gene, so we looked for other sources of ribosomal frameshift data to test the accuracy of PRFect. Xu et. al. examined the conservation of the translational frameshift in bacteriophage tail assembly chaperone genes and found 28 phage genomes with two genes that share homology to the two parts of the tail assembly chaperone gene of phage lambda. Out of the 28 *joined* TAC genes of the Xu phages, ten were identified correctly as utilizing ribosomal frameshifting, and only one gene (a hypothetical one) was incorrectly predicted to contain a frameshift.

### The Frame Shift Database

The third source of frameshift data was the FSDB, which was a comprehensive compilation of experimentally known or computationally predicted data about programmed ribosomal frameshifting. The database contains 253 frameshifts from all domains of life and functions; unfortunately, it has not been updated since its inception fifteen years ago.

### Covid19

The last data source for our testing sets was the single genome of SARS-CoV-2, the virus that causes Covid 19. The genome contains 12 genes, one of which is a polyprotein that contains a ribosomal frameshift to translate a much longer version of the polyprotein. PRfect exactly predicts the single frameshift in the polyprotein gene, and without any other False Positive predictions of the other genes.

## Discussion

### SEAPHAGES

Genomic sequencing is at the forefront of most biological research, partly due to the ever-increasing accessibility of sequencing technology. Phage and prophage genomes are being increasingly studied for their roles in almost every facet of human life; from health, to industry, to environment. The SEA-PHAGE initiative is one of many numerous sequencing efforts underway to help better our understanding of the full breadth of genomic diversity and molecular complexities of the bacteriophage world. The function of PRFect is to detect those translationally abnormal genes in phage genomes that are potentially subject to ribosomal programmed frameshifting. As a side benefit, PRFect also uncovered previously unknown “shifty” sequence motifs that are likely to induce ribosomal slippage and promote frameshifting. One of the issues when working with the SEA-PHAGES genome database, like most sequence databases in general, is that it has a lot of misannotated data. There are no estimates on how prevalent tail assembly chaperone frameshifting is; while it is somewhat conserved among double-stranded tailed phages, it is certainly not present in every single SEA-PHAGE genome. Out of the 3,679 SEA-PHAGE genomes we downloaded, only 2,476 contained a *joined* PRF gene (2,397 tail assembly chaperones and 79 hypotheticals). Of the remaining 1,203 SEA-PHAGE genomes without an annotated PRF gene, a fuzzy word search revealed that 347 had a single *tail assembly chaperone* (TAC) gene, while 342 had two TAC genes. It would be reasonable to presume that a genome with two TAC genes should accordingly have a *joined* PRF gene, so we took the “False Positive” genes (those predicted by PRFect to be joined but that were not joined in the file) at each of the five validation split levels, and grouped them into very general functional categories and a spillover OTHER category (the exact numbers and mapping is found in the supplemental documents). As usual, the genes of *hypothetical* function eclipsed all other categories. The second largest functional category was *tail assembly chaperones*, suggesting that many SEA-PHAGE genomes lacking a *joined* PRF gene were due to errors caused by incomplete annotations. Around 80% of these “false positive” TACs were from genes that were not *joined* in the genome file, while the remaining 20% were from TAC genes that were *joined* in the sequence file, but were considered as *false* due to the location of the predicted frameshift being more than 10 bp away from where it was annotated as occurring in the sequence file. It is unknown whether the PRFect prediction is wrong or whether the genome annotation location is wrong. Of the 161 TAC genes where PRFect predicted a different location for the frameshift, there were 136 that did not have any motif at the original location, suggesting that perhaps the genome annotation is wrong. We suspect that many of the genes annotated as encoding *hypothetical* proteins are also TAC genes, since the evidence for adding a *joined* PRF gene to the genome annotation is the presence of two adjacent TAC genes. If the genome is so divergent that one or both of the TAC genes lack sufficient sequence similarity to known TAC genes, they will be reported as *hypothetical*, and subsequently will not be *joined* during the manual annotation of the genome.

**Figure 6).**
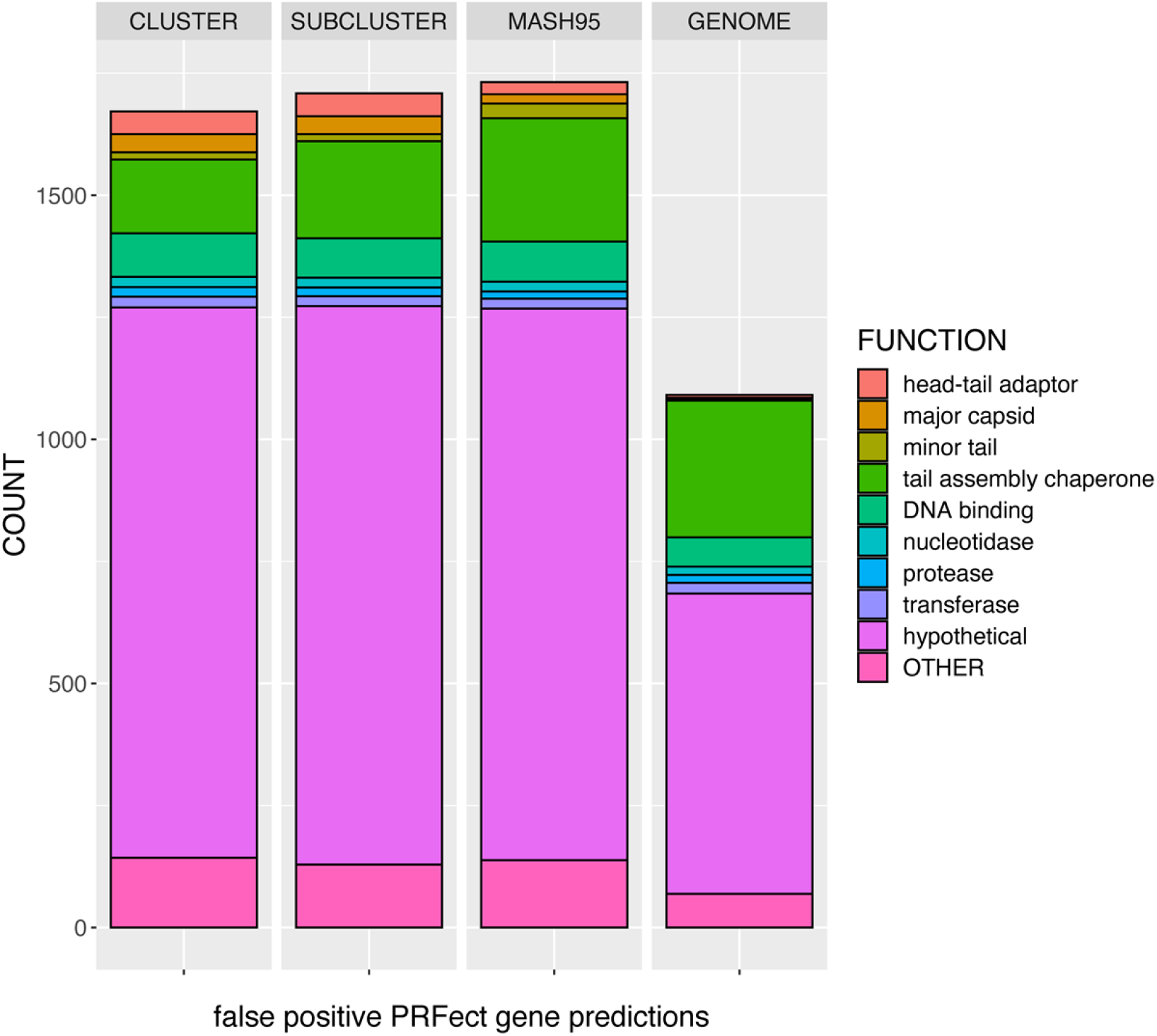
The various gene functions of the false positive PRF predictions during the leave-one-out validation levels. False Positive genes were two adjacent genes predicted by PRFect to contain a frameshift, but that were not joined accordingly. Gene functions were grouped into 12 broad categories based on fuzzy word matching and manual group estimation to simplify the data. Some categories were groupings of similar cellular functions, like the various enzymes, while some were just standardizing the existing function names. For example, there were 25 different spellings of hypothetical protein and 29 different versions of tail assembly chaperone. Gene functions that could not be reasonably assigned to one of the 12 categories were grouped into the OTHER category. The counts are twice the numbers shown in the previous figure because if a gene is predicted by PRFect to be joined when it is not actually a joined gene, it will thus be two genes of potentially different functions.

Specific protein functional annotations allowed us to perform a literature search for evidence supporting the true- and false-positive classifications. For example, the gene encoding *Cro* is known to contain a frameshift when expressed in *E. coli* (33). Other genes in the “False Positives” that are known to contain a frameshift were: *lysins*, *methyltransferases*, *DNA/RNA polymerases*, *IS3 family transposases*, *RusA*/*RuvC*, and *major/minor capsids* (34–41); *minor tail* and *tape measure* genes which could be incorrectly labeled *TAC* genes; as well as *VIP2 ADP-ribosyltransferases* and *MuF-like* which have been found in genomes as a single fusion gene (42). The relevance of fusion genes is that some T4-like phages do not utilize ribosomal frameshifting to get the two forms (short and long) of the TAC gene but instead carry one copy of the short gene and one copy of the longer gene that is a fusion of the short and longer downstream part (43). Therefore, if two genes occur in some genomes as a single fusion gene, there is the possibility that when those genes occur as separate genes in different frames, they may utilize PRF to get the fusion protein during translation.

One last source of concern with the SEA-PHAGES data is that there were 103 genomes that did not have a slippery site motif within 10 bp of the annotated putative frameshift site. This could be due to the location being wrong, sequencing error, or the slippery site is not one of the 10 motifs that PRFect looks for. Of those 103 genomes, there were 22 that did not have a slippery site motif anywhere within the overlap region, which suggests that either the slippery sequence is of a motif that PRFect does not look for or that the sequence contains sequencing error.

### The RECODE data

One of the more critical cellular properties that characterize forward frameshifts is the ribosome encountering a rare codon, which PRFect cannot adequately determine for the RECODE data because it comprises partial genomes. An alternate mechanism that can also induce forward frameshifts is that instead of a rare codon, the ribosome pauses at a *stop* codon. PRFect treats all of the 64 possible codons equally to calculate the frequency of each codon in the genome, and since stop codons are one per gene, they can appear as rare codons. PRFect still requires that a *three* or *four* motif be present before the *stop* codon, and only 33 of the forward frameshift slippery sites in the RECODE data have such a motif. Of the remaining 448 forward RECODE frameshifts that use a different motif, 295 have the nucleotides `CUU`, and 139 have the nucleotides `UCC` just before the *stop* codon. In the first case of `CUU_U` (the fourth base must be `U`, since all three *stop* codons start with `U`), is perfectly base paired with its cognate tRNA `GAA`, and the ability of G to weakly bind `U`, allows for the `GAA` to pair with `UU_U` in the +1 frame (44). All of the +1 RECODE frameshifts with the `CUU*` motif (the asterisk denotes a stop codon) belong to bacterial release factor 2 (*prfB*) genes, which suggests that they comprise a negative feedback loop to finely tune the level of PrfB protein in the cell. When cellular levels of PrfB are high, it binds to the ribosome complex and terminates translation at the `CUU*` stop codon, resulting in translation of the shorter nonfunctional PrfB protein variant. When cellular levels of PrfB are low, the ribosome encounters the slippery site and pauses much longer since there is no PrfB to terminate translation. Eventually, a forward base slip occurs that shifts the ribosome into the +1 frame, forming the longer functional PrfB protein. When randomizing the codon before the stop, it was shown that the next two most slippery motifs are `CCC` and `UUU`; which supports our hypothesis that both *three* and *four* are valid slippery site motifs for +1 forward frameshifts. In the second case of `UCC_U`, no explanation for the codon:anticodon re-pairing is provided in the literature, and the `AGG` tRNA has only two base positions that pair (the second and *wobble* third via G:U pairing) with the +1 codon `CCU.` Interestingly, the gene that is translated by this frameshift is ornithine decarboxylase antizyme (OAZ1), which inhibits polyamine synthesis and import. The +1 frameshift is more frequent at high cellular polyamine levels, leading to more OAZ1 protein, which reduces polyamine levels. One way this is accomplished is that during the translation of OAZ1 in the absence of polyamines, the nascent peptide interacts with the ribosome and prevents its own synthesis, leading to increased polyamine levels (45). Polyamines are also known to bind to rRNA (ribosomes), mRNA, and tRNA; therefore, polyamines may alter the translation machinery and make a transient motif out of the `UCC` nucleotide pattern.

Another possibility is that they are not +1 frameshifts at all; it was shown *in vitro* that the mammalian OAZ1 frameshift is ostensibly a +1 shift (8), though strangely when the exact same sequence was expressed in *S. cerevisiae*, proteomics revealed that the frameshift is reached through a -2 shift rather than a +1 shift (46). The OAZ1 frameshift also has a pseudoknot 3’ of the slippery site, which is usually utilized in backward frameshifts since the downstream secondary structure impedes the forward progress of the ribosome. It is possible that the pseudoknot is only one part of bidirectional PRF control and that polyamines influence the stability of the pseudoknot as another aspect of the negative feedback loop control. If we were to add `CUU*` and UCC* to the motifs that are searched for, as well as use the entire genome, presumably PRFect would detect the *prfB* and OAZ1 frameshifts correctly (which make up 58% of the RECODE data), and the recall would go up dramatically for the RECODE dataset.

### The Xu phages

We had expected that PRFect would perform quite well on the Xu dataset since it was comprised of the same TAC gene used for training. However, PRFect correctly identified 11 of the 28 frameshifted TAC genes from the phage genomes. The cause of three of the missed frameshifts was that the slippery sequence motif at the frameshift sites was not one of those searched for by PRFect. The remaining incorrect predictions seem to be caused by a combination of the GC content being much lower or the distance of the slippery site (N) from the in-frame stop codon being much higher (>30 nt) than the SEA-PHAGES data. The GC content has been shown to affect the minimum free energy of the downstream secondary structure (Supp Fig 2). The average GC content of the SEA-PHAGES used for the training was 64%, compared to only 47% in the Xu phages. The slippery site’s average distance (N) from the in-frame stop codon is 26 nt in the training set and 36 nt in the Xu dataset.

### The FSDB data

Due to the size of the FSDB entries, many of which are full prokaryotic genomes or entire Eukaryotic chromosomes full of thousands of genes, we were unable to manually curate the data to ensure that both the slippery site indicated in the FSDB was present and add it if it was not, or to remove those *joined* genes that were not the indicted frameshift. Although there were only 253 frameshifts listed on the FSDB website, there were 897 total *joined* coding sequences across all of the genomic files. The FSDB data was included as a test set for completeness and illustrative purposes, with the precision and recall performance of PRFect being marginal at best but the result does show the performance accuracy of PRFect. The accuracy is based on the *false-positive* to *true-negative* ratio, and considering that there were 150,000 non-*joined* genes in the dataset and that almost all were accurately predicted as true-negatives, PRFect had an accuracy of >99% (Supp Fig 3).

### Atkins

The Atkins database consists of eukaryotic viruses that tend to organize their genes into a single gene spanning the entire genome as one uninterrupted open reading frame (31). The gene is translated as one large polyprotein which is then later cleaved by proteases into the constituent proteins. Many of the frameshifts were shifts into a frame that contained an early stop codon, thus acting as a method to create a much shorter version of the polyprotein. Consequently, the length of the slippery site from the *in-frame stop* codon (N) was the entire length of the second frameshifted portion of the gene. The average N for the SEAPHAGES data that was trained upon was 26 nt, while the average N for the Atkins genomes was 169 nt. The length N is one of the features that was added to help discriminate the True Positive slippery sites near to the *in-frame stop* codon from the *true-negative* slippery sites occurring much farther in the 5’ direction of the overlap. Of the 43 genomes with N greater than 51 nt, only two were successfully predicted as containing a frameshift by PRFect, and oddly enough, they are -2 frameshifts that present as +1 frameshifts with the slippery site motif of *three* nucleotides in a row (GUU_UUU).

One of the concerns that we have deferred until now that applies to all ribosomal frameshift analysis is that there are more than just -1 (backward) and +1 (forward) frameshifts. Various translational coding anomalies can cause the ribosome to skip around on the mRNA more than just a single base, including -2 shifts, +2 slips, +5 steps, +6 hops, and even a colossal +50 bp jump (5,47). Despite the range of nucleotides that may be jumped, all frameshifts are still present in the data as either -1 or +1 offset. Because there are only three coding frames per strand direction, so if there is a shift from the 0 frame, regardless of direction, you are either in the -1 or +1 frame. A +2 shift presents the same as a -1 shift, a -2 shift presents the same as a +1 shift, a +4 shift presents the same as a +1 shift, and so on. There is also the possibility of a shift that would put the ribosome back into the 0 frame, i.e. via a -3 or +3 shift. The functional purpose of this would be to skip one or more codons, however there are tRNA reassignments that allow *st*op codon readthrough (essentially a +3 frameshift), while there are no documented cases of ribosomal programmed frameshifts of multiples of 3 nucleotides.

#### SARS-CoV-2

Unlike many of the Atkins viral genomes with PRFs that cause a shorter version of the polyprotein to be translated, the PRF in SARS-Cov-2 frameshift causes a longer version of the polyprotein to be translated. Thus, the distance of the slippery site from the in-frame stop codon is only 15 nt. This short distance is just one of the contributing properties that enables PRFect to perfectly predict the frameshift in the polyprotein gene without any other False Positive predictions on the other genes. This single example shows that despite PRFect being trained on one specific coding gene of prokaryotic phage genomes, it uses universal cellular properties rather than sequence homology. The generalized model allows it to identify frameshifts in a broad range of coding genes and diverse taxonomical clades.

#### Comparing Performance

PRFect is the first and only tool that predicts programmed ribosomal frameshifts. The only other tools available, FSFinder and KnotInFrame, do not predict PRFs but show all possible locations of a given slippery site motif within a genome (11,12). FSFinder finds four motifs: *threethree* for the backward PRFs; and *threeStop*, *UCCstop*, and *CCUstop* for the forward frameshifts. All these motifs appear relatively frequently in a genome by chance, so a second version of the tool (FSFinder2) added the requirement that the motif is located within a gene overlap region (Figure 3) to help reduce the *false-positive* rate, but even this more restrictive version still has very low *precision* when run on our datasets (Supp Fig 6). FSFinder had the best *precision* (41%) on the RECODE data composed of single genes and the worst *precision* (1%) on the FSDB composed of huge eukaryotic genomes. The recall was not much better: it did find the slippery site of the single frameshifted gene in SARS-Cov-2 but only found around 25%-56% of the slippery site in the frameshifted genes of the other datasets. KnotInFrame looks for only the *threethree* motif with downstream secondary structure, then scores and sorts the results showing however many the user chooses, with a default of 11 (per strand). The performance was also unacceptable: KnotinFrame did find the slippery site in the single frameshifted gene of SARS-Cov-2, but it also reported 21 other matches for the *threethree* motif in the genome. The real design flaw of KnotInFrame was evident in the SEA-PHAGES data, which had 3,479 genomes with 2,476 annotated *joined* genes, where KnotInFrame found 523 of the true-positive slippery sites and more than 48,000 extra (*false-positive*) slippery sites locations (Supp Fig 7).

#### Future Improvements

We did not consider taking into account the specific nucleotides that can occur in the base positions of a motif during the development of PRFect. The original XXXYYYZ heptamer motif has a consensus that only specific nucleotides can occupy each position: where X can be any nucleotide, Y can be A|U, and Z can be A|U|C (48). For the SEAPHAGES data, we observed no such positional base limitations, and each of the four nucleotides was found in each of the three XYZ positions of the true-positive joined genes with *threethree* motifs (Supp Fig 5). Other motifs had more narrow nucleotide limitations, such as *fivetwo* which had the nucleotides GGGGGAA across all of the thousand motifs of the SEAPHAGES data. The problem with implementing positional nucleotide constraints is that our SEAPHAGES training data is biased toward high GC% content and highly repetitive, which would cause machine learning models trained on the data to follow the bias. One aspect that could prove to be an improvement to the motifs used by PRFect is considering G:U base pairing of the slippery site codons and the bound tRNAs in the E and P sites. In the previously mentioned *fivetwo* observed motif of GGGGGAA, the second tRNA CUU rebinds with the codon GGA. Only the first position (and the wobble third) are complementary using regular A:U and C:G pairing. However, if G:U base pairing is considered, then all positions are complementary. It may be that some of the motifs that we hypothesize to exist are instead limited to only with specific nucleotides in certain motif positions that allow for G:U base pairing. An example is that the most common bases observed for the hypothetical *threetwotwo* motif (CCCGGAA) have G:U pairing that mimics a *threethree* motif. Though supporting our postulate of new alternative motifs is the fact that the majority of the *twoonefour* motifs cannot be explained through G:U pairing alone.

Another possible improvement would be to adjust the codon rarity based on identical codons upstream of the potential frameshift site codon to consider the temporal nature of the tRNA pool. If a codon is repeated once or more in short succession within a coding gene, that tRNA can be consumed quicker than tRNA is recharged. Thus, two or more moderately *infrequent* codons repeated back-to-back would deplete their cognate tRNA from the pool and ultimately act as a rare “hungry” codon. Tandem repeats have been shown to induce ribosomal frameshifting *in-situ* in *E. coli* (49–51), are responsible for the frameshifting associated with many human diseases (52,53), as well as translational-pausing involved in the synthesis of amino acids and polyamines (54,55). Dividing the waiting A-site codon frequency by the number of times it occurs within some upstream window of a given length would give a more nuanced measure of how “hungry” a rare codon is. Additionally, adjusting the frequency of a given codon by its near-cognates could also improve the model even further.

## Conclusion

PRFect is the only tool currently available for predicting programmed ribosomal frameshifts in a given genome with a very high degree of accuracy. It is easily installable with a single command, and since the code is open source, we have established a path towards improving PRFect by including additional cellular properties, such as new motifs. We expect PRFect to change and adapt to the field as more is discovered about programmed ribosomal frameshifting and as ever-increasing genomic data is added to the model. For PRFect to predict that two adjacent genes within a genome annotation are one frameshifted gene with its two (or more) constituent parts split into different frames, those parts need to be predicted as distinct genes by gene calling software. The four most popular gene-finding tools, GeneMark, Glimmer, Prodigal, and Phanotate, all function by looking for *start* codon to *stop* codon pairs within the same contiguous frame of the genome (22,56–58). The downstream second part of a ribosomally frameshifted gene does not necessarily have a start codon, so traditional gene prediction algorithms might not find it. However, the codons that can serve as *start* codons (AUG, GUG, and UUG) also occur quite frequently within a gene, where they code for standard amino acids, which can allow for gene finders to call the downstream fragment as a gene. When the second part of a frameshifted gene does not have a valid start codon, we have a new gene finding tool, Genotate, that detects coding regions within a genome without relying on *start* codon to *stop* codon pairs (in preparation). So rather than relying on other third-party gene finding tools that may not detect all the fragments of a ribosomally frameshifted gene, we will have in place the means to processes the nucleotide genome with Genotate in order to get the coding regions, which are then given to PRFect so that it may predict programmed ribosomal frameshifts between adjacent coding regions.

## Supplemental Figures

**Supplementary Figure 1).**
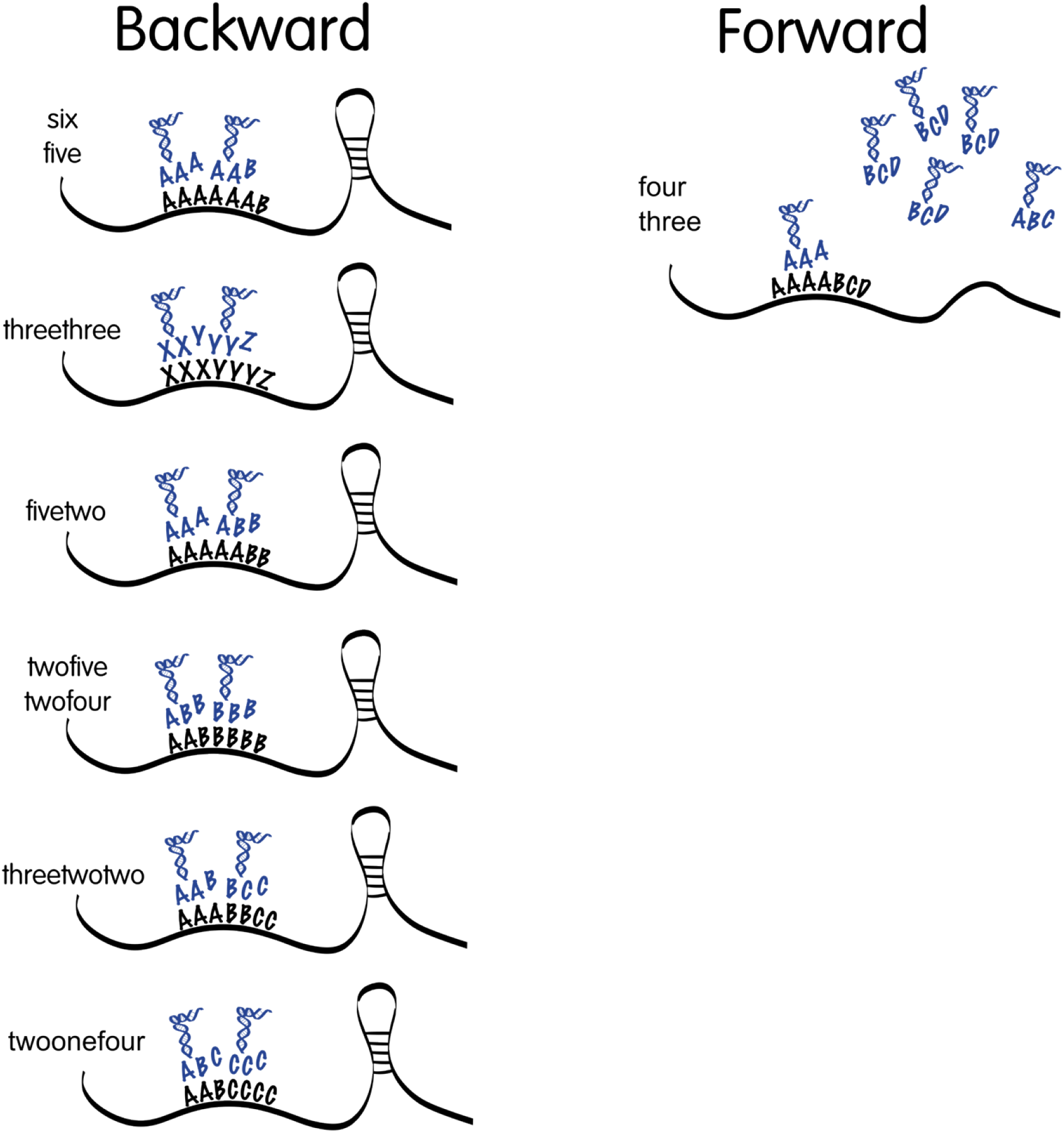
The ten different motifs that PRFect uses as slippery sites to then predict which of those sites find are involved in programmed ribosomal frameshifting. Shown is the originally proposed XXXYYYZ motif for backward frameshifting, along with 9 other proposed motifs that might also allow for the wobble base pairing mismatch initialy discovered with the XXXYYYZ motif. The motifs are named according to the repeated bases (i.e. six is 6 bases in a row), are shown in descending order based on the probabilty of seeing that motif by chance alone, and some are grouped together for brevity. The black letter correspond to the bases (codons) of the mRNA transcript while the blue base correspond to the bases (anti-codons) of the two bound tRNA. The tRNA are shown in their frameshifted position (backwards or forwards) and show how the third weak wobble position of the tRNA might not match the new frameshifted pairing. For example in the original threethree motif the tRNAs XXY and YYZ are bound to the mRNA and then shift back one base so that only their 1^st^ an d 2^nd^ bases are paired correctly while the 3^rd^ wobble bases are mismatched. The probabilty of seeing three (and four) bases in a row in given genome is quite common, so for the forward frameshifts we also require that the codon of the waiting A-site (ABC) is rarer than the codon of the +1 A-site (BCD). The codon rarities are calcualted at run time on the input genome by iterating through all of its coding genes and counting the occurrence of each of the 64 different codon posibilties. Requiring the +1 A-site to be more common than the waiting +0 A-site codon in forward frameshifts helps to cut down on false-positive preditions but mainly serves to speed up runtimes; since three bases in a row is quite common and calculting the MFE of a window is computational time intensive.

**Supplementary Figure 2.**
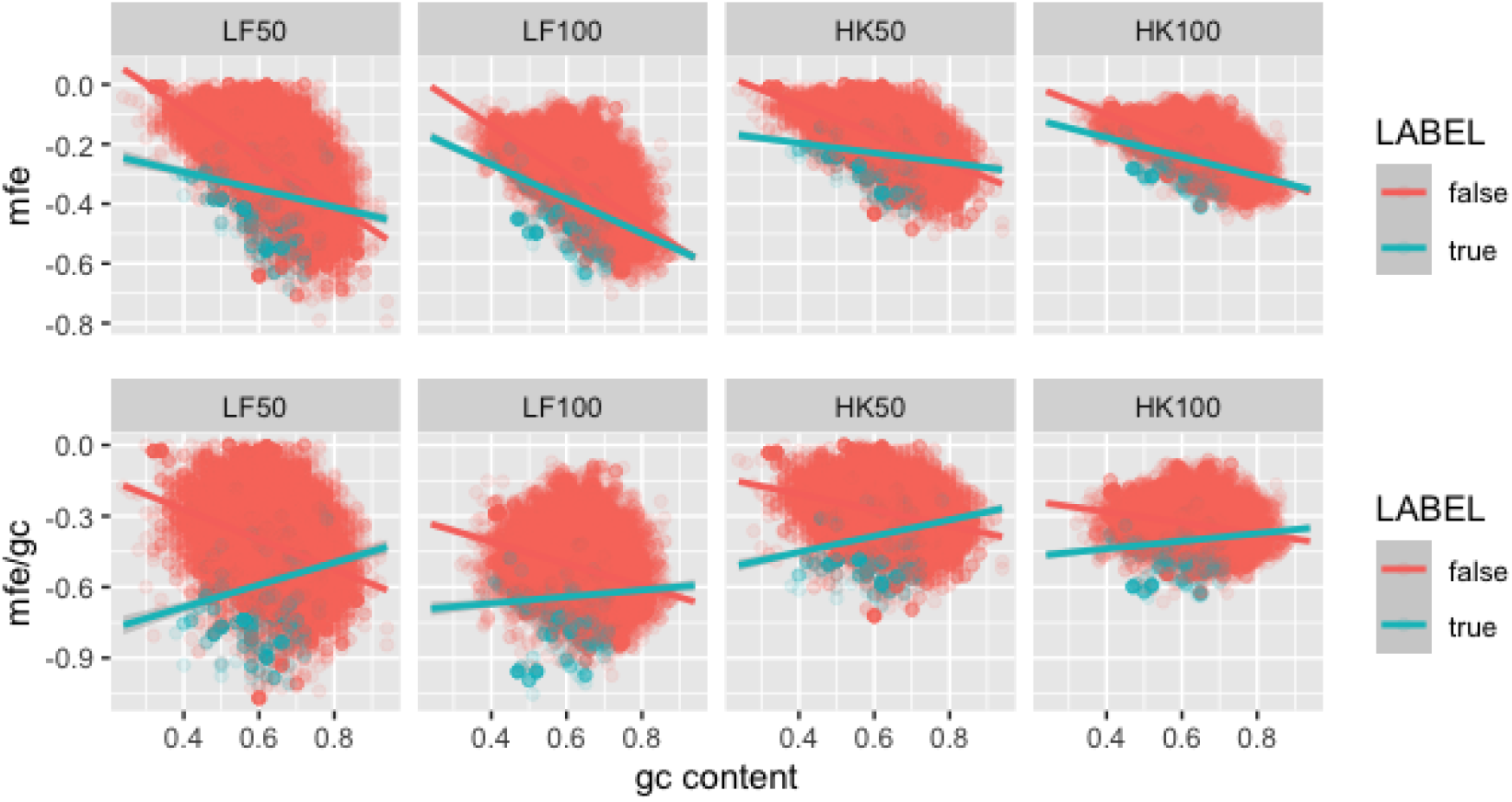
The programs LinearFold (LF) and HotKnots (HK) were used to calculate the minimum free energy (MFE) for 50bp and 100bp windows downstream of slippery sites in the SEAPHAGES data from *joined* genes (*true*) known to frameshift and from genes that are adjacent and not *joined* in the annotation file (*false*). The top show the linear realtionship between the MFE and the GC% content thaat would impair a model trained on data biased towards high or low GC content genomes. The genomes of the SEAPHAGES data tend to be higher in GC content, so in order to help PRFect perform on genomes of any GC% content, we divide the MFE by the GC to help normalize the MFE to the GC.

**Supplementary Figure 3.**
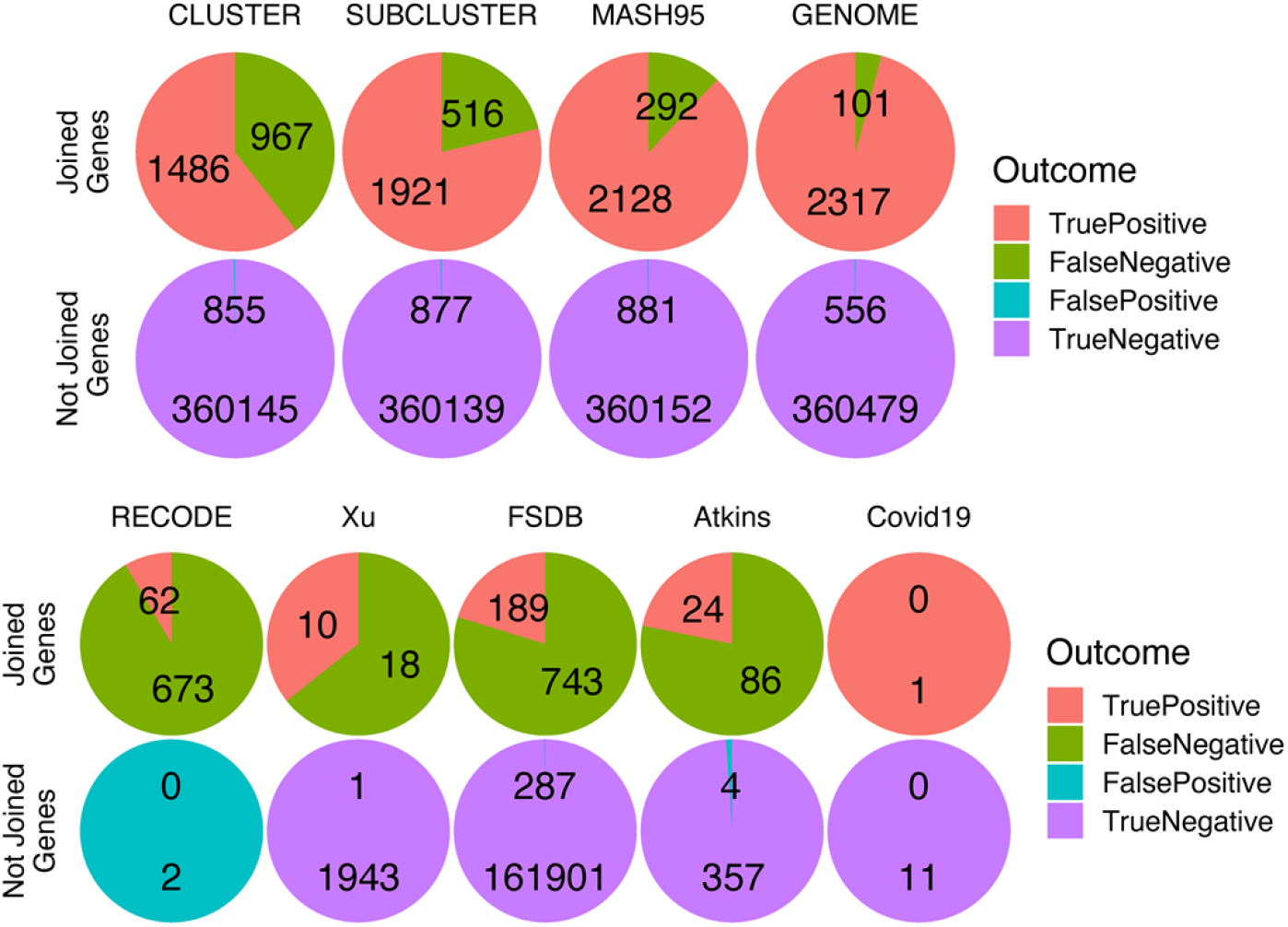
The full results of PRFect on the various datasets

**Supplementary Figure 4).**
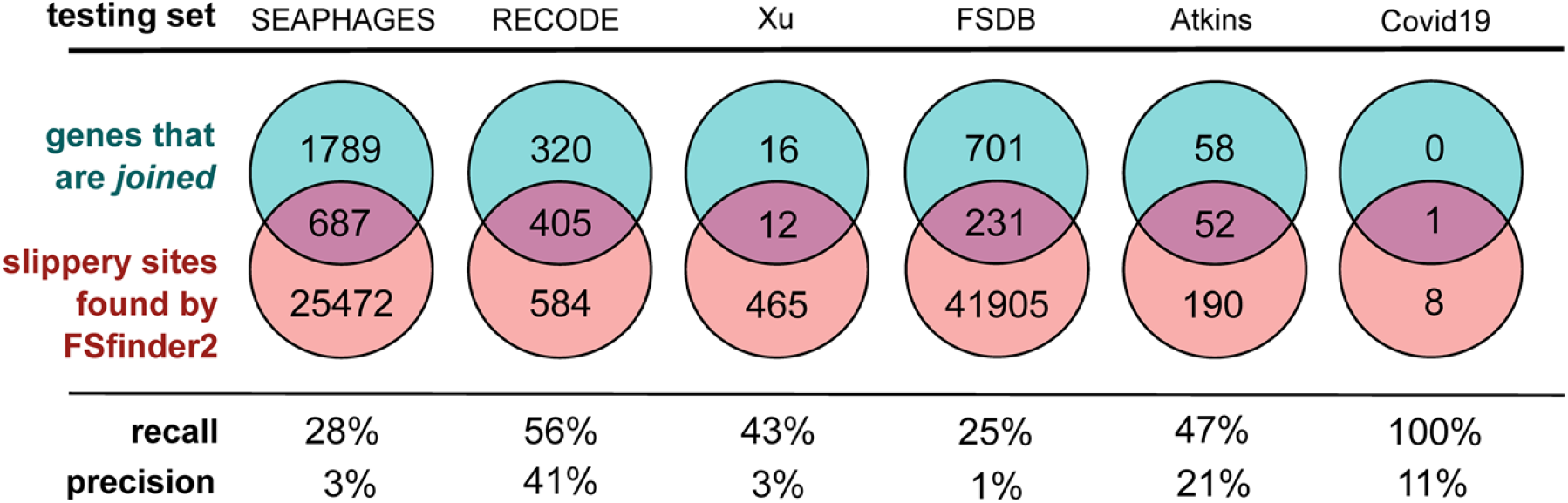
The performance of FSFinder2 on the various datasets.

**Supplementary Figure 5).**
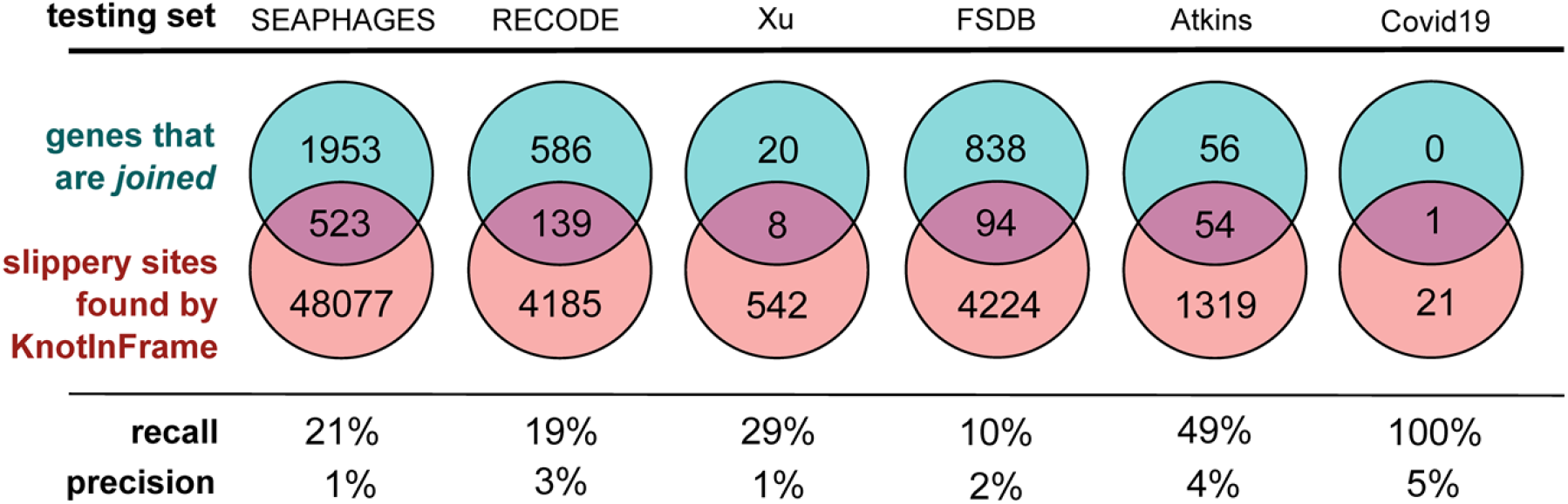
The performance of KnotInFrame on the various datasets.

**Supplementary Figure 6).**
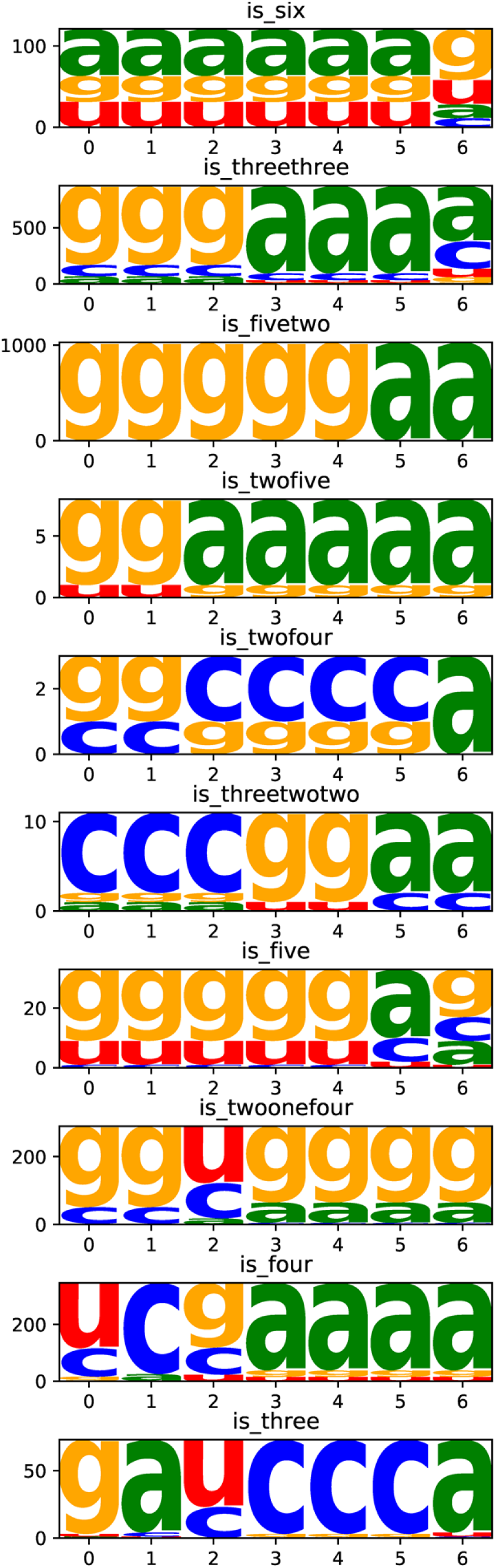
The nucleotide frequency at each base location in the various motifs of the true-positive slippery sites

## Notes

### Competing Interest Statement

The authors have declared no competing interest.

